# Ranking genome-wide correlation measurements improves microarray and RNA-seq based global and targeted co-expression networks

**DOI:** 10.1101/299909

**Authors:** Franziska Liesecke, Dimitri Daudu, Rodolphe Dugé de Bernonville, Sébastien Besseau, Marc Clastre, Vincent Courdavault, Johan-Owen de Craene, Joel Crèche, Nathalie Giglioli-Guivarc’h, Gaëlle Glévarec, Olivier Pichon, Thomas Dugé de Bernonville

## Abstract

Co-expression networks are essential tools to infer biological associations between gene products and predict gene annotation. Global networks can be analyzed at the transcriptome wide scale or after querying them with a set of guide genes to capture the transcriptional landscape of a given pathway in a process named Pathway Level Correlation (PLC). A critical step in network construction remains the definition of gene co-expression. In the present work, we compared how Pearson Correlation Coefficient (PCC), Spearman Correlation Coefficient (SCC), their respective ranked values (Highest Reciprocal Rank (HRR)), Mutual Information (MI) and Partial Correlations (PC) performed on global networks and PLCs. This evaluation was conducted on the model plant *Arabidopsis thaliana* using microarray and differently pre-processed RNA-seq datasets. We particularly evaluated how dataset x distance measurement combinations performed in 5 PLCs corresponding to 4 well described plant metabolic pathways (phenylpropanoid, carbohydrate, fatty acid and terpene metabolisms) and the cytokinin signaling pathway. Our present work highlights how PCC ranked with HRR is better suited for global network construction and PLC with microarray and RNA-seq data than other distance methods, especially to cluster genes in partitions similar to biological subpathways.

## Introduction

Constructing global gene co-expression networks is a popular approach to highlight transcriptional relationships (edges) between genes (vertices). The ‘Guilt-by-Association’ (GBA) principle supposes that genes sharing similar functions are preferentially connected and aims at predicting new functions for proteins by determining how their respective encoding genes are co-expressed with others using a reference dataset containing known gene functions such as the Gene Ontology (GO)^1^. Defining edges connecting genes remains a critical step in global co-expression network construction. Expression data (microarray or RNA-seq) are used to construct expression matrices (genes x samples) and to calculate a distance or a similarity for each possible gene pair. The resulting pairwise distance matrix is then thresholded to obtain an adjacency matrix that discriminates relevant edges. Only edges with a distance below (or a similarity above) the set threshold are considered significant and retained for network construction. The procedure is expected to remove non biologically relevant gene associations while retaining the relevant ones and can be assessed with any reference dataset. Alternatively, guide gene sets may be used to extract more human-readable information from large networks in a process named Pathway-Level Correlation (PLC)^2–6^. This approach aims at capturing the best transcriptional associations of a gene set and at highlighting functional gene groups such as known subpathways in this set. There are two types of approaches to determine transcriptional associations of genes: those that are supervised and those that are unsupervised. Supervised approaches such as regression and machine learning based methods require a prior knowledge which is used as a training dataset to recover biologically relevant gene associations. The superiority of supervised methods in extracting potential physical regulatory interactions between genes has been demonstrated using simulated and real *E. coli* and *S. cerevisiae* subnetworks^7^. This study has revealed that prediction accuracy is higher with smaller networks and concluded that inferring genome-scale networks remains elusive unless performing a feature selection step to reduce inference problem size (because of the under determined nature of current expression datasets). Among the unsupervised methods, four are commonly used and have been thoroughly tested. The first approach is Mutual Information (MI) which measures a statistical dependence between two variables^7^. It is based on density function estimates and has been shown to perform well with non linear relationships^8^. The second approach which relies on integrating multiple transcriptional associations is Partial Correlation (PC). PCs are generally calculated from multiple linear regression and include a variable selection step^9^. PCs aim at explaining a gene’s expression profile by a small number of strongly correlated genes after eliminating those less correlated that do not significantly explain this gene’s expression profile. The two last methods are Correlation Coefficients (CCs), either Pearson CC (PCC) or Spearman CC (SCC), which are the classical estimators of linear transcriptional relationship among genes^9, 10^. CCs are 2-dimensional distance measurements because a CC between two genes does not take into account the expression of the remaining transcripts in the whole transcriptome. To compensate for this lack, these approaches have been improved by using ranked CCs instead of raw values. Ranking CC implies that for every gene, all CCs calculated with the N-1 remaining genes (where N is the number of genes) are ranked from 1 to N. Within a pair of genes A and B, rank(A to B) differs from rank(B to A) because the two genes display different expression profiles and different relationships with the remaining transcripts in the transcriptome. Two related ranking methods have been developed. One is mutual ranking (MR, geometric mean of the two ranks) which has been shown to improve GO term recovery with PCC using large microarray data from Arabidopsis, Human, mouse and rat^11^. MR has been successfully used in multispecies analysis of co-expression modules^12^. Another is Highest Reciprocal Ranking (HRR, maximum value of the two ranks)^13^. MR and HRR are thought to be more integrative than unranked CCs because they depend on other CC values around that of a gene pair. Although not as robust as supervised methods, unsupervised methods can efficiently capture relevant gene associations as previously shown^8^. These authors have shown that non parametric CC and MI calculations were more efficient than PCC on a small dataset. Among other unsupervised methods, SCC calculations have been similarly shown to outperform other distance measurements in Human expression data^14^. In this case, SCC were calculated from RNA-seq or microarray data in order to construct several smaller networks subsequently aggregated to yield the final network. We firmly believe that genome-scale networks inferred with CCs, especially when combined with a ranking procedure, are helpful to find new associations between genes. Although CCs are not efficient in detecting non linear associations^8^, gene-to-gene relationships have been predicted to be essentially linear^15^ suggesting that CCs are valuable distance measurements. To date, there is no clear evaluation of how ranked CCs affect genome-scale network reconstruction with RNA-seq data in comparison with other unsupervised methods. We evaluated ranked CC, raw CC, MI, and PC performance in global and targeted network construction using Arabidopsis microarray and differentially processed RNA-seq expression data (Figure 1). Performance was measured as network ability to capture biologically relevant gene associations found in a Gene Ontology (GO) annotation reference set but also to correctly cluster guide genes in PLC. Global network quality was first evaluated according to the different dataset x distance measurement combinations. The resulting global networks were next interrogated in PLC analyses with five different guide gene sets corresponding to four different metabolic pathways and one signaling pathway. Whereas metabolic pathways have relatively clearly defined and partially linear partitions, signaling pathways usually involve post transcriptional regulations and a more intricate organization, which might render gene transcriptional associations less evident. We looked at the dataset x distance measurement combinations optimizing pathway reconstruction and maximizing co-occurrence quality between microarray and RNA-seq networks. Our results show that, of the six methods evaluated, PCC ranked with HRR generated the best biologically relevant networks according to initial guide gene representation and clustering in distinct modules. In addition, it offers the possibility to merge subgraphs obtained by microarrays and RNA-seq to generate high confidence networks.

**Figure 1.**
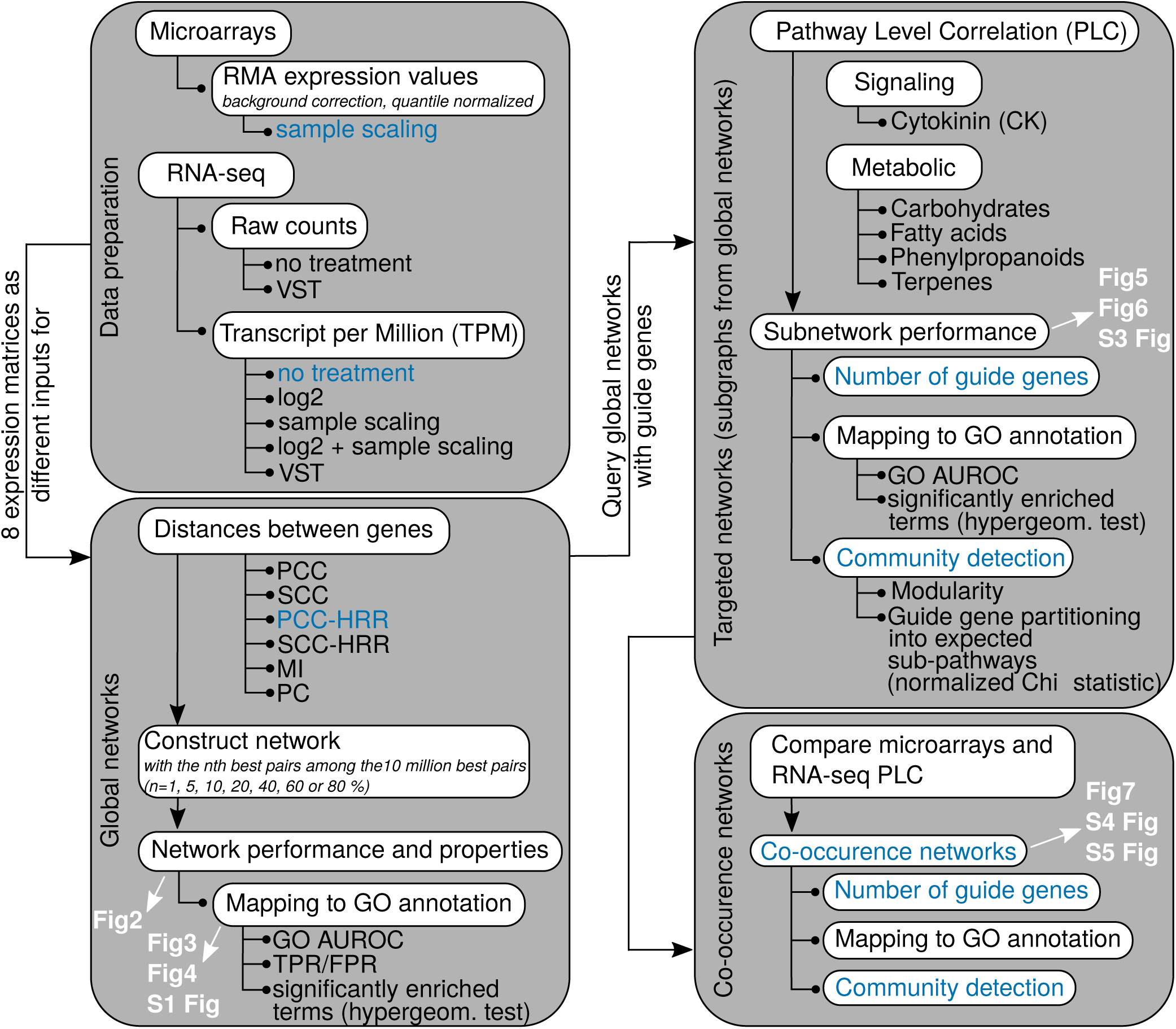
Workflow for global and targeted network analyses. One microarray dataset and a RNA-seq dataset prepared according to 7 normalization procedures were used to generate eight expression matrices analyzed with six different distance measurements (Pearson’s or Spearman’s Correlation Coefficient, unranked or ranked with HRR, Mutual Information (MI) or Partial Correlations (PC)) to obtain 48 distance matrices. Each of these matrices was thresholded to obtain global networks at different confidence thresholds. Global networks were evaluated and also queried with specific guide gene sets reflecting 5 different pathways in a process named Pathway Level Correlation (PLC). The resulting subnetworks were evaluated and used to construct co-ocurrence networks between microarray and RNA-seq datasets. In white are indicated the figures corresponding to the different steps analyzed. Dataset x distance combinations are indicated in blue and characteristics that are improved by these combinations.

## Results

### Inferring global co-expression networks and comparing correlation measurements

Large co-expression networks were obtained by varying the confidence threshold (correlation value above or rank below) within lists containing the 10 million best gene pairs from eight different datasets and six data measurement combinations (Figure 1). Each of the 10 million best pair lists was filtered at different confidence thresholds (1, 5, 10, 20, 40, 60 or 80% best pairs from these lists) to evaluate the effect of network size on performance. Expression datasets included a microarray-based expression matrix and seven RNA-seq based expression matrices normalized with different methods to evaluate their effect on network inference: transcript per Million (TPM), log2 TPM, sample scaled (ss) TPM, ss log2 TPM, raw counts, variance stabilized transformed (VST) raw counts and VST-TPM. The six distance measurements were: raw PCC, raw SCC, PCC-HRR, SCC-HRR, PC and MI. Each network performance was considered as a network ability to capture edges corresponding to functional associations found in the GO reference dataset and was evaluated in 4 different ways (Figure 2): GO term enrichment (GO terms that are significantly enriched with gene pairs from the co-expression network), a ROC curve constructed with TPR and FPR calculated for each confidence threshold and two ROC analyses based on the GBA concept, an average 3-fold cross validated neighbor voting (NV) AUROC and a global AUROC. AUROCs correspond to Area Under Receiver Operating Characteristic curves calculated for every network either from each GO (with three test sets obtained after hiding part of the gene labels, NV AUROC corresponding to the average of AUROCs for all GO terms) or the whole annotation dataset (global AUROC). AUROCs are used as global indicators of a dataset performance, a value of 0.5 indicating a random attribution of labels in the network and a value of 1 indicating a perfect match with the reference dataset. AUROC*>*0.6 may be considered as moderate^14^. In global TPR vs FPR curves, the line extending from (0,0) to (1,1) has an AUROC=0.5 and points above this line indicate more predictive networks than a random selection (Figure 2). The GO annotation table was filtered to perform these analyses by removing weakly represented or non-specific GO terms (*>*5 or *<*100 genes).

**Figure 2.**
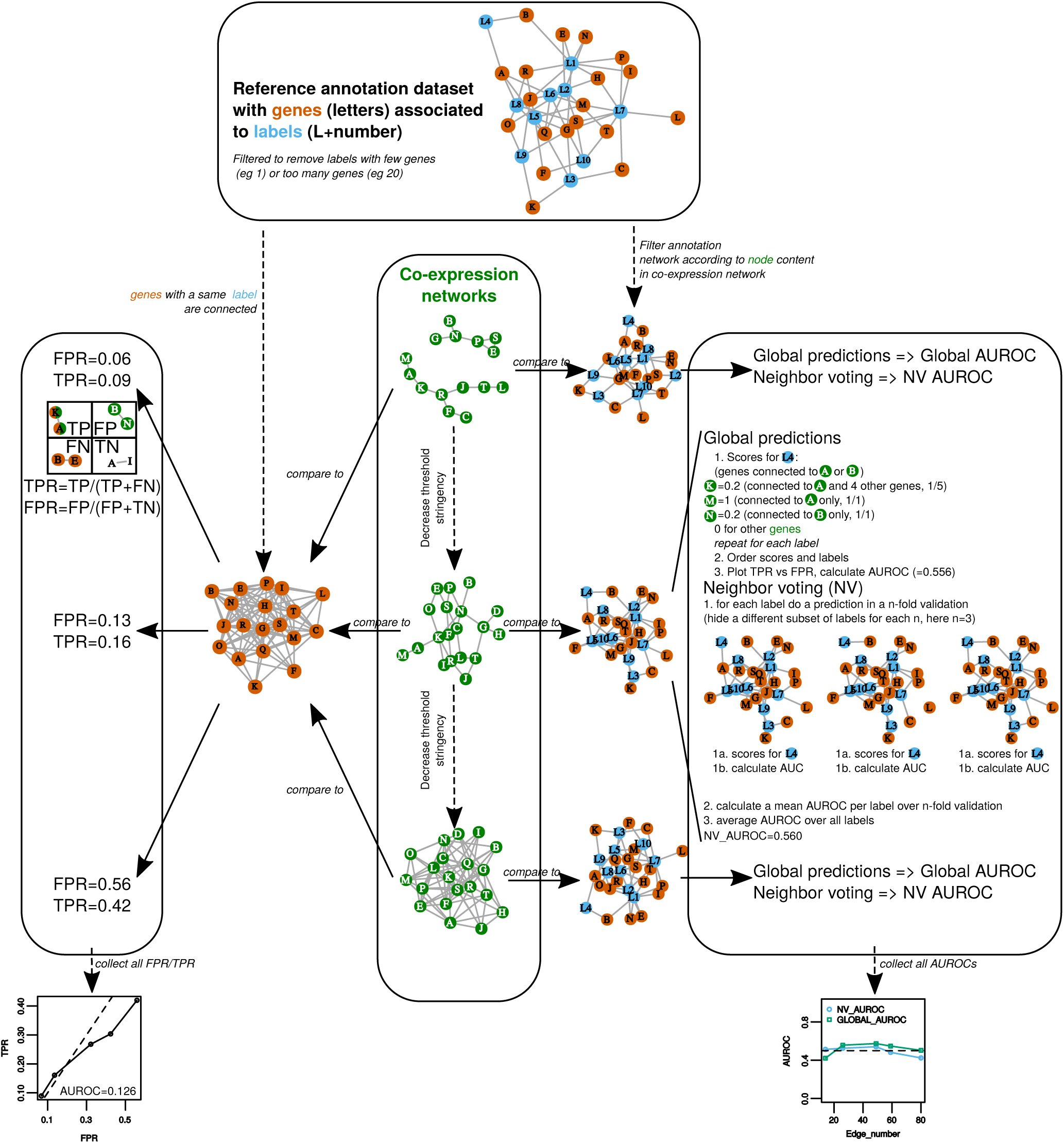
Network performance. This small example describe strategies to evaluate networks according to a reference functional annotation. Co-expression networks were obtained for each dataset x distance measurement combination (Figure 1) at different confidence thresholds, resulting in networks increasing in size with lower stringency. A total evaluation was made with True Positive Rate (TPR) vs False Positive Rate (FPR) analysis (left panel) by classifying edges as True positives (TP), False Positives (FP), False Negatives (FN) or True Negatives (TN). Single network evaluation was performed by calculating AUROCs with the EGAD R package, either as a global prediction or using a neighbor voting (NV) algorithm with a 3-fold cross validation (right panel). All indicated values are in accordance with the small networks in this example. In addition to these 3 evaluations (FPR vs TPR, global AUROC and NV AUROC), GO term significant enrichment was statistically tested with a hypergeometric distribution (not shown in this example).

Figure 3 displays TPM network evaluation at different confidence thresholds and Figure 4 shows networks having 1 million of edges across all dataset x distance combinations. Metrics for all other dataset x distance measurement combinations are presented in Supplementary Figure 1 online. All networks combined, pairwise correlations between enriched GO counts, global and NV AUROC performance metrics were moderate (Spearman’s *rho>*0.4) but significant (*p<*0.001) indicating these three performance metrics evaluated networks in different ways. The highest correlation was observed between NV AUROC and enriched GO counts (*rho*=0.70, *p<*0.001) showing their consistency. The NV AUROC was the most positively correlated with edge number (*rho*=0.55, *p<*0.001) suggesting that decreasing the confidence threshold and adding more edges in networks did not result in a significant increase in false positives. This was confirmed by the partial ROC curves (obtained for a maximum FPR at 10 million edges) drawn from the TPR and FPR (Figure 3, Supplementary Figure 1 online), where up to 10 million best pairs, TPR increased faster than FPR. Although counts of significantly enriched GO terms were positively correlated to NV AUROC, we observed a slight decline in the largest networks which might reveal a saturation in these enriched GO terms. It is possible that with the hypergeometric testing, some GO classes are fully enriched in smaller networks leading to a decrease in their significance as network size increases. The global AUROC displayed a very low variation (min=0.55, average=0.61, max=0.68) and was significantly correlated to vertex number (*rho*=0.43, *p<*0.001) only. This observation suggests that the global AUROC is not an appropriated measure in our case.

**Figure 3.**
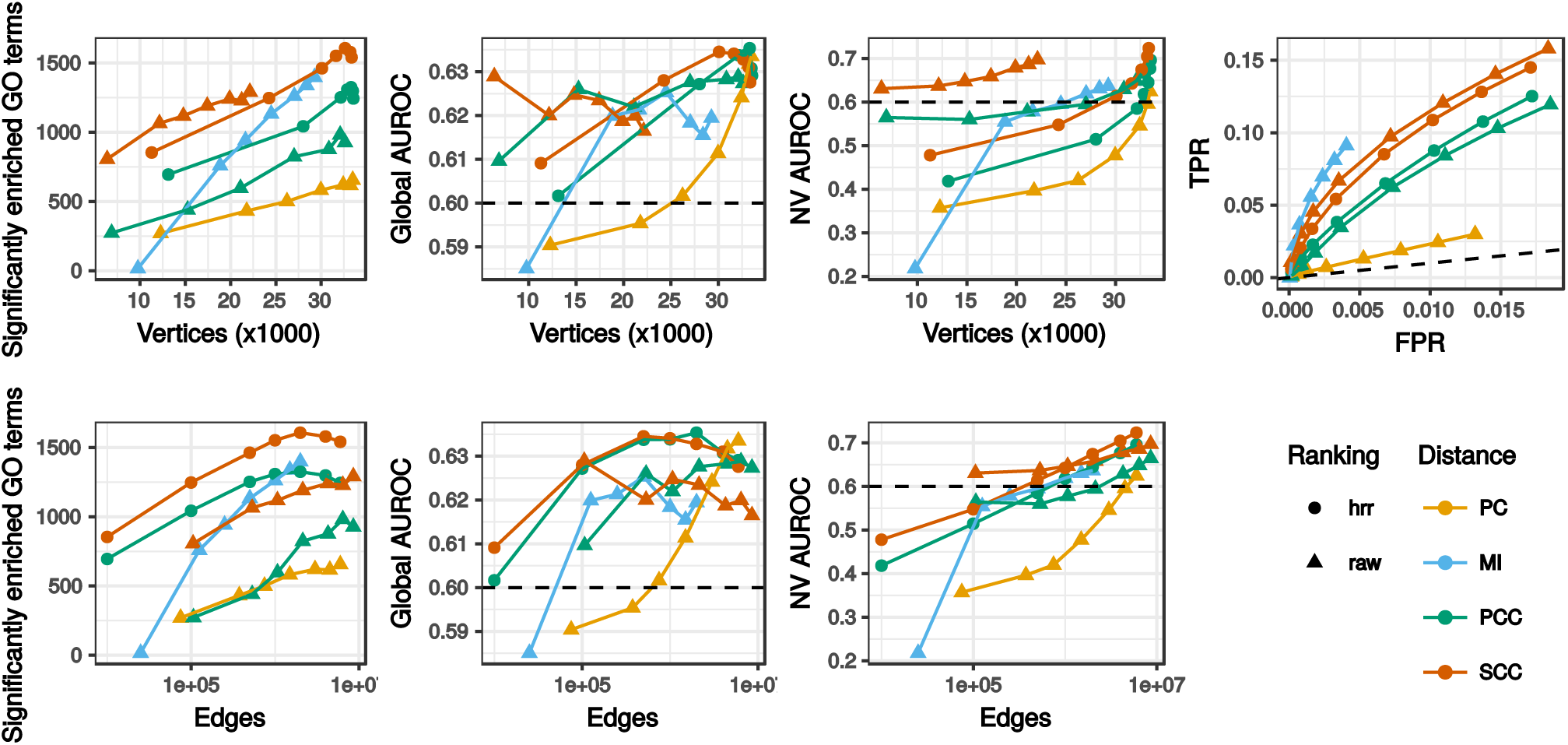
Global network characteristics. Only results for the RNA-seq TPM dataset without further normalization are shown. The horizontal dashed line indicates a 0.6 AUROC value taken as a threshold separating good and poor network predictability. In the TPR=f(FPR) panel, the dashed line corresponds to a random selection (with AUROC*<*0.5). This panel is partial and the highest FPRs correspond to 10 million gene pairs.

**Figure 4.**
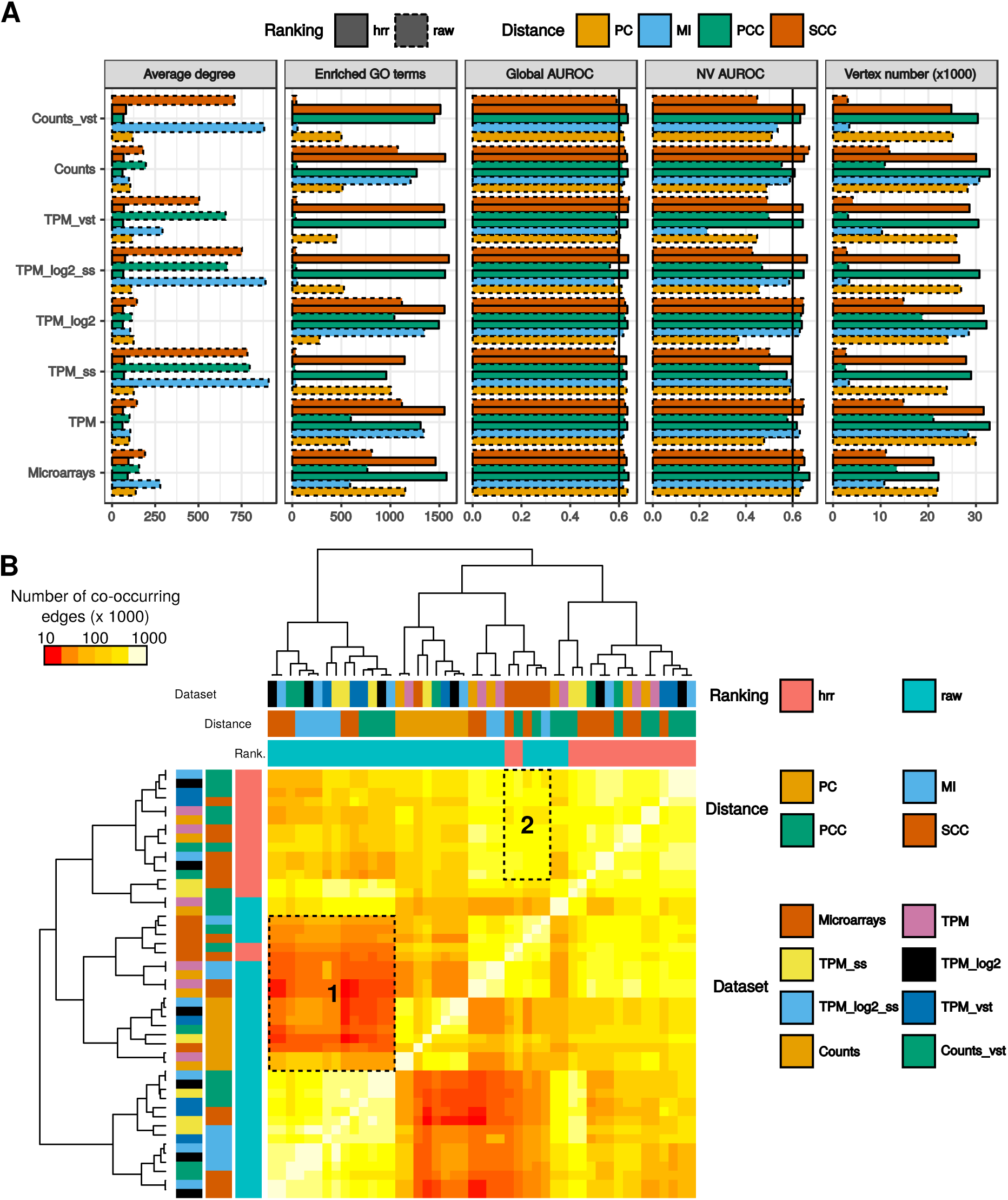
Comparison of dataset x distance measurement combinations for networks with a million gene pairs. Network topology and performance in GO recovery were analyzed (A). Vertical lines at 0.6 indicate AUROCs above which network predictability can be considered as moderate. Co-occurring edges were also counted in every possible comparison between 2 networks (B). Area 1 corresponds to RNA-seq networks having few genes in common with PC networks and microarrays networks and area 2 to combinations maximizing edge co-occurrence between microarray and RNA-seq.

At equivalent edge numbers, different distance measurements generated networks varying considerably in vertex number (Figure 4A, Supplementary Figure 1 online). Considering all datasets and distance measurements, raw PCC, raw SCC and MI resulted on average in fewer vertices and higher node degree (vertex number/node degree: 13,164/511, 9,986/465 and 14,074/468 respectively) than PCC-HRR, SCC-HRR or PC (26,645/116, 24,731/124 and 23,966/166 respectively). This trend was clearly observed when setting an edge number to 1 million (Figure 4A). Expression networks constructed from microarrays, TPM, TPM log2, and counts displayed very similar ROC curves: PC based networks followed random predictions (NV AUROC=0.5) and the other distance measurements were above the random prediction with similar AUC (Supplementary Figure 1 online). This was confirmed for PC by NV AUROC and enriched GO term counts. Performance of the other distance measurements in the global TPR/FPR curves did not exactly match that measured with AUROCs. Taking the TPM dataset as an illustration (Figure 3), the MI ROC curve was above the others while NV AUROC for similar edge numbers was slightly below that measured for SCC. This was probably due to differences in network topologies (see above) and the procedures underpinning the two evaluations. The global TPR/FPR curve does not measure a network predictability *per se* as NV AUROC does and considering any gene pair sharing a same GO term as valid could have overestimated TP (Figure 2). As a general trend, raw PCC and raw SCC generated smaller networks than PCC-HRR and SCC-HRR but displayed similar TPR/FPR curves, *i.e*. for a similar performance, HRR-ranked CC networks had more vertices and fewer edges than raw CC based networks (Figure 3). CC ranked with HRR always generated relevant networks for TPM ss, TPM log2 ss, TPM VST and counts VST, which was not the case for raw CC (Supplementary Figure 1 online). These normalizations induced strong biases in CC distribution as revealed by thresholds used to obtain the 10 million best pairs (Supplementary Table 1 online) but these biases were compensated by HRR. Taken together, these results revealed that HRR CCs are able to generate complete genome-wide networks with good performances similar to other classical measures such a MI and PC. Node degree AUROC measures whether genes are more likely associated according to their number of connections rather than to their function. A positive correlation was found between NV AUROC and degree AUROC (*rho*=0.47, *p<*2e-16) indicating that highly predictive networks (NV AUROC*>*0.7) also had a higher node degree AUROC. Node degree AUROC was generally under 0.55. We therefore considered that in our conditions, this bias was only limited. Concerning edge co-occurrence between the different dataset x distance combinations, the lowest conservation was observed with raw (MI, PCC and SCC) RNA-seq datasets and PC networks and microarrays networks (Figure 4B, area 1). More co-occurring edges were found when microarray networks were compared to RNA-seq networks obtained with CC-HRR (mean of 97,646 vs 25,277; Figure 4B, area2). This indicated that microarrays and RNA-seq networks were more comparable when obtained with HRR, reinforcing their validity. The previous section focused on global network properties. Community detection procedures can be applied to such global networks to cluster tightly connected genes into modules. In our case, we rather used a knowledge-driven approach known as Pathway-Level Correlation (PLC) to extract gene pairs associated within a given pathway (Supplementary Figure 2 online). PLC are particularly interesting in plants for example to decipher incomplete specialized metabolic pathways. It aims at capturing a transcriptional landscape for genes known to be involved in a given pathway, in order to highlight their organization as well as finding new genes (transporters, transcription factors,…) associated with the process. In the next part, we evaluated the ability of all previous networks to capture relevant information associated with four metabolic and one signaling pathways. We selected two primary metabolic pathways (carbohydrate and fatty acid metabolisms), two specialized (secondary) pathways (phenylpropanoid and terpenoid metabolisms) and the cytokinin signaling pathway.

### Assessing PLC quality: trade-off between GO term representation and guide genes

The PLC procedure is expected to cluster together guide genes with many co-expressed genes (‘associated genes’) and to reflect the subpathway organization (Figure 5A). For PLC, we systematically removed all genes showing a degree value of 1 (*i.e*., those connected to only one guide gene). However we included edges between associated genes if they were found among edges retained at the selected threshold. Using five pathways (Table 1, Figure 5B, Supplementary Table 2 and Supplementary Figure 3 online), we extracted five PLC from the global networks generated above to determine the best suitable dataset x distance measurement combinations. All pathways have modular structures with gene sets forming specific sub-pathways (also called partitions or modules). We expected that PLC would be able to reconstruct such a partitioning, by connecting guide genes with associated genes. The phenylpropanoid pathway contains a core module composed of 3 genes leading to a precursor used by 3 other distinct subpathways^16–19^(Figure 5B, Table 1, Supplementary Table 2 online). The three other metabolic pathways, carbohydrates, fatty acids and terpenoids, were structured in modules as described on the KEGG database^20^ (Table 1, Supplementary Table 2 online). The fatty acid pathway contains 97 genes divided into 6 modules. The central carbohydrate metabolism contains 202 genes partitioned in 8 modules. Finally, the terpene pathway has 64 genes partitioned into 6 modules. Pathway organizations were used as indicated in the KEGG database (apart from phenylpropanoid pathway which was manually curated from our previous work) and compared to PLC subnetworks. The plant cytokinin (CK) pathway is known to regulate many processes in plant physiology and is hierarchically organized in three levels: a histidine kinase receptor, a transducer (histidine phosphotransfer proteins) and a response regulator (type A/B/C) which may act as a transcription factor^21^(Table 1, Supplementary Table 2 online). Although CK pathway members are relatively well known, each level is represented by several members which may have specific roles and it is still unclear how they biologically interact with each other to drive a specific physiological response. We expected that PLC would group some of these actors according to specific physiological responses. CK pathway includes both transcription activating and repressing activities (via response regulators) and post-transcriptional (phosphorylations) and would therefore be an excellent test of PLC applicability on associations expected to be more complex than in metabolic pathways. In addition, we included other histidine kinases integrating other signals and known to crosstalk with the CK pathway^22^. We therefore included 2 ethylene receptors, ETR1 and ERS1 to determine whether they could be clustered with CK histidine kinase. The initial pathway was not partitioned into sub-pathways but rather into 5 levels (receptor, transducer, type A/B/C response regulator) because interactions between specific actors of each level are not completely understood.

**Figure 5.**
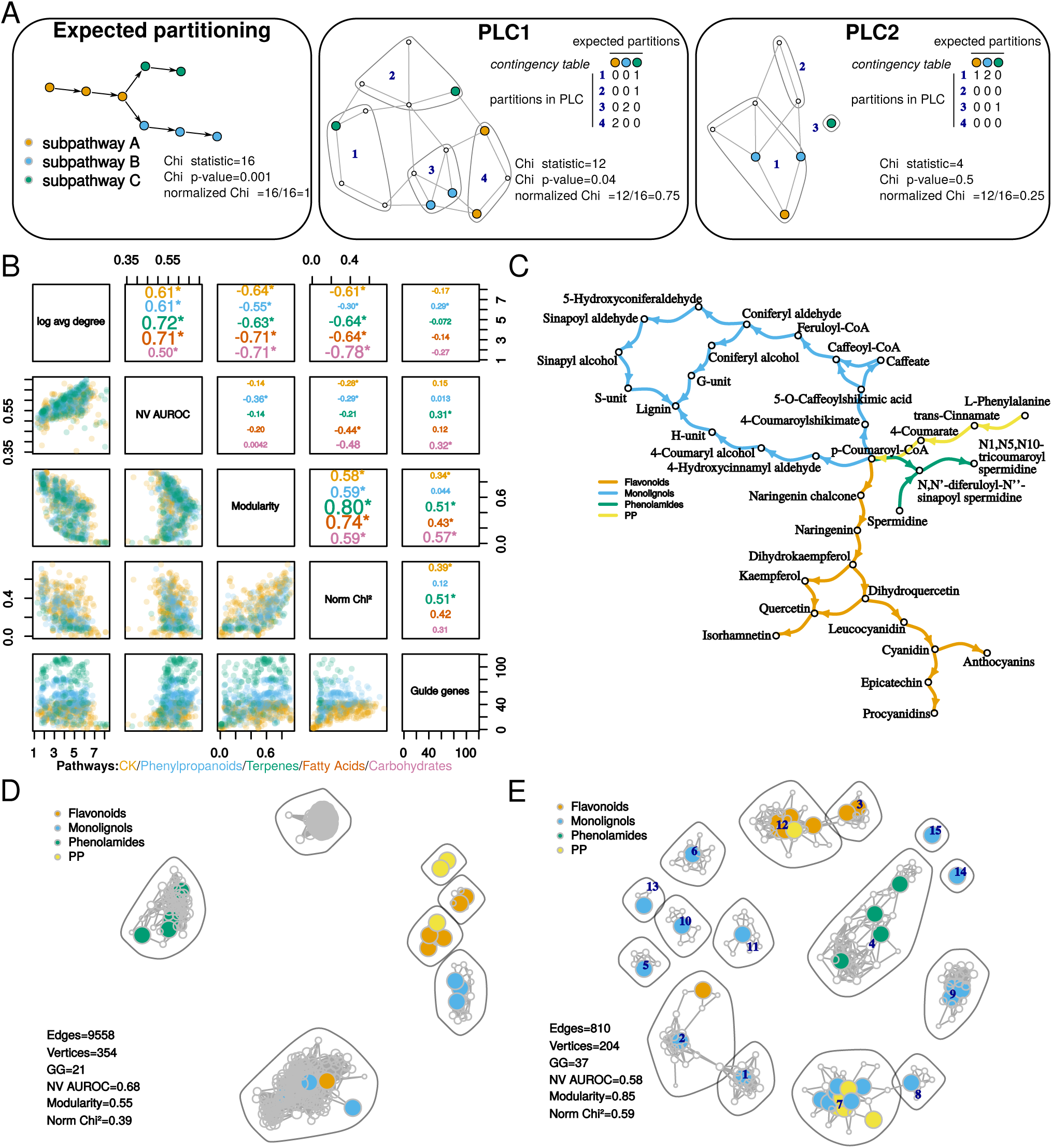
Trade-off in PLC subnetworks between performance in GO term recovery and partitioning guide genes into expected communities. (A) Example showing normalized Chi^2^ statistic and *p*-value calculations comparing guide gene distribution into PLC communities (numbers in deep blue within polygons) to the expect partitioning (left; 3 subpathways). Two PLCs (one with a good partitioning (center); one with a weak partitioning (right)) are shown here but the contingency matrix used in Chi^2^ calculations is described for only one of them (center). (B) Pair plot showing correlations (Spearman’s rho, asterisks show significance *p<*0.001, upper panel) and scatterplots (lower panel) between average network node degree, NV AUROC, normalized Chi^2^, modularity and the number of guide genes in the network. Each point in the lower panels (scatterplots) represent one network for which 2 characteristics (eg NV AUROC and modularity) are compared. Data are presented for each pathway separately with a specific color. (C) The expected partitioning of phenylpropanoid related guide genes was compared to two PLC: (D) higher predictability and lower modularity (microarrays raw PCC) and (E) lower predictability and higher modularity (microarrays PCC-HRR). In D and E, colored vertices correspond to genes encoding enzymes catalyzing steps of similar color in C. Community (surrounded by grey polygons) numbers in E are indicated in deep blue and can be used to access Supplementary Table 3 online.

**Table 1.** Pathway description. ? Indicates that partition in sub-pathway is not known.

| Pathway | Genes | Number of subpathways | Subpathway names (KEGG module accession) |
| --- | --- | --- | --- |
| Phenylpropanoids | 43 | 4 | core phenylpropanoid (PP), flavonoids, monolignols, phenolamides |
| Fatty acid | 97 | 6 | fatty acid biosynthesis (initiation (M00082), elongation (M00083), its ER-localized part (M00415)), jasmonic acid phytohormone biosynthesis (M00113) and β-oxidation (M00086 and M00087) |
| Carbohydrate | 202 | 8 | glycolysis (Embden-Meyerhof pathway (M00001) and the core module involving three-carbon compounds (M00002)), neoglucogenesis (M00003), pyruvate oxidation (M00307), citrate cycle (M00010), pentose phosphate pathway (M00004, M00006 and M00007) |
| Terpenes | 64 | 6 | mevalonate (M00095), methylerythritol (M00096), C10-C20 isoprenoid (M00366), beta-carotene (M00097), abscisic acid hormone (M00372) and phytosterol (M00371) biosynthetic blocks |
| Cytokinin signaling | 37 | ? | ? |

Subgraphs of global networks were constructed for each pathway by retrieving edges involving at least one guide gene and were partitioned into communities with a fast greedy algorithm designed to maximize network modularity and which has been shown to extract relevant communities from large networks^23^. We compared guide gene distribution in these communities to target subpathways using a normalized Chi^2^ test which values range from 0 to 1, 1 being the expected partition and 0 a random partition of guide genes or very few guide genes (Figure 5A). All networks having a Chi^2^ *p*-value*>*0.05 were considered to have a Chi^2^ statistic equal to 0. PLC performance in recovering GO terms was evaluated by counting significantly enriched GO terms and by calculating a NV AUROC for each network. A good PLC was expected to contain a large number of guide genes and to have both a good score in grouping them into expected partitions (high normalized Chi^2^ value) and a good score in overall biologically relevant edge recovery (NV AUROC*>*0.6). We first analyzed correlations between all these metrics (NV AUROC, number of guide genes and Chi^2^ statistic) together with two topological metrics (mean node degree and modularity), for each pathway separately (Figure 5B). Strongest correlations were observed between NV AUROC and mean node degree (*rho>*0.5, *p*e*<*0.001) and between modularity and normalized Chi^2^ (*rho>*0.59, *p<*0.001). We found that PLC performance (NV AUROC) was almost negatively correlated with normalized Chi^2^ (*rho<*-0.2) indicating that guide genes were clustered correctly at the expense of capturing GO associated gene pairs. Given the CK pathway structure, partitioning based on protein functions (receptor, transducer or response regulator) did not resulted in high Chi^2^ values, suggesting that partitions in the co-expression networks contained guide genes from different levels, reinforcing the existence of specific sub-pathways. These results indicated a trade-off in PLC between edge quality and guide gene partitioning. A visual examination of PLC with either lower modularity and higher NV AUROC (Figure 5D) or higher modularity and lower NV AUROC (Figure 5E) revealed that PLC with higher modularity as well as higher Chi^2^ values displayed a biologically relevant organization. Such subgraphs had generally a lower average node degree and a higher representation of guide genes rendering their analysis more convenient. Taking the phenylpropanoid pathway as an example, the PCC-HRR based TPM network (Figure 5E, with a higher modularity) correctly clustered genes from the core phenylpropanoid (PP) and the flavonoid modules while the raw PCC network did not (Figure 5D, with a higher NV AUROC). Similar results were observed with the four other pathways with either microarray or RNA-seq datasets (Supplementary Figure 3 online). Modularity and normalized Chi^2^ could therefore be considered as consistent quality metrics for PLC. NV AUROC should also be considered to ensure that subgraphs had a minimum predictability (*>*0.55).

### HRR-CCs optimize recovery and clustering of guide genes in PLC

The best performing dataset x distance measurement combinations were searched by analyzing NV AUROC, modularity and normalized Chi^2^ among networks with a Chi^2^ *p<*0.05. Statistical effects of dataset, distance, ranking and their interactions on subgraph characteristics were analyzed by ANOVA for each pathway. Ranking and distance measurements had generally the strongest effects on modularity and normalized Chi^2^ (*p<*2e-5) (Figure 6A & B). Ranking had a significant effect on NV AUROC (*p<*0.01) but was weaker than distance measurement (*p<*1e-4). Datasets only had a significant effect on modularity (*p<*0.002). Significant interactions were rarely observed between these three factors (*i.e*. in few pathways and with a weak effect). This revealed that the different RNA-seq normalizations had only minor effects on these PLCs. Taken as a whole, networks obtained with raw datasets had a significant higher NV AUROC (t-test, mean in raw=0.58, mean in HRR=0.57, *p<*0.01) but significant lower modularity (mean in raw=0.35, mean in HRR=0.68, *p<*2.2e-16) and lower normalized Chi^2^ value (mean in raw=0.20, mean in HRR=0.35, *p<*2.2e-16) (Figure 6A). It therefore appeared that clustering guide genes correctly was improved with CC ranked with HRR at the expense of performance. NV AUROCs in HRR-based networks were generally higher than 0.55, indicating an average low performance in GO capture (Figure 6A). In non-ranked distances, PC resulted in the weakest NV AUROC, while MI and raw SCC based networks displayed the highest NV AUROC (Figure 6B). This weakness in PC based networks was compensated neither by a higher modularity nor by a higher normalized Chi^2^ statistic.

**Figure 6.**
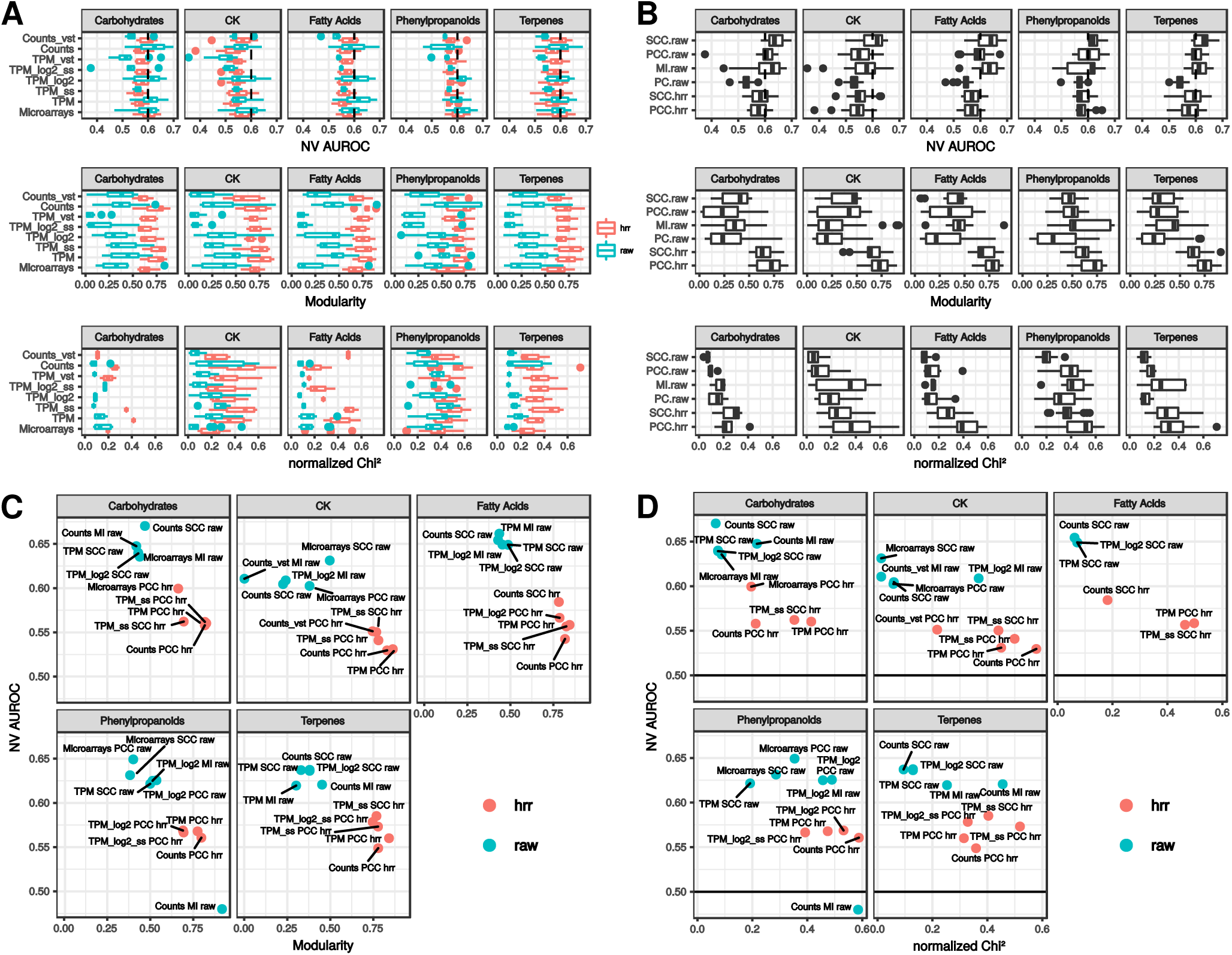
PLC subnetwork performance. Performance in capturing GO terms (NV AUROC), modularity and normalized Chi^2^ value distribution in interactions between datasets and ranking methods (A) and between distance measurement and ranking methods (B) showing the dominant effect of the ranking procedure (raw vs HRR) on these metrics. (C) Modularity and NV AUROC of the five top NV AUROC networks and 5 top modularity networks. (D) Normalized Chi^2^ statistic and NV AUROC for the same networks.

A more detailed examination of best PLC subgraphs maximizing either modularity or NV AUROC, revealed that each of the five pathways involved specific dataset x distance measurement combinations. PCC-HRR based networks were always found to maximize modularity (Figure 6C) and normalized Chi^2^ (Figure 6D) with almost all datasets. Raw distance based PLCs had a higher NV AUROC and some of them also had a good modularity but they also had a lower normalized Chi^2^ statistic indicating they contained fewer guide genes (*e.g*. raw RNA-seq counts with raw SCC in the terpene PLC). The results suggest that PCC-HRR could be used as a reliable distance measurement whatever the dataset. Careful analysis of PLC obtained from PCC-HRR revealed the presence of relevant associations in each PLC (Supplementary Figure 3 and Supplementary Table 3 online). For example, community 12 from the phenylpropanoid PLC obtained with microarray data processes with PCC-HRR (Figure 5D) contained AT1G06000 encoding a Flavonol 7-O-rhamnosyltransferase and was clearly associated with other genes from the flavonoid sub-pathway. This gene was not detected in the raw PCC PLC (Figure 5C). Other examples are highlighted in yellow in Supplementary Table 3 online.

### Vertex and edge co-occurence in microarray and RNA-seq based PLC subgraphs

Edge co-occurrence in networks constructed from expression datasets obtained by different technologies may be considered as a further validation. Quantifying gene expression with microarrays relies on probe hybridization by sequence complementary while with RNA-seq, short reads are mapped back *in silico* to the reference transcriptome. The two main differences between these technologies are (i) the number of quantified transcripts (due to the completion of genome annotation) and (ii) the dynamic range (fluorescent probe intensities for microarrays, *in silico* read counts for RNA-seq). Because microarrays and RNA-seq technologies differ, edges co-occurring in networks obtained from these two technologies are probably more relevant. In Figure 4C, we analyzed co-occurrence in global networks and found that HRR ranked CCs apparently increased the number of co-occurring edges between microarrays and RNA-seq. To get more insights into co-occurrence in PLCs, common edges and vertices were counted in pairwise intersections of networks (RNA-seq vs microarrays) obtained with the six distance measurements and set at a 1,000 vertices. The resulting intersection networks were further characterized by the number of represented guide genes, their normalized Chi^2^ statistic, modularity and NV AUROC. This evaluation was performed with the RNA-seq dataset expressed as TPM only because we showed in the previous section that normalization methods had a minor impact on PLC. In addition, TPM networks with raw distance methods had enough vertices to correctly extract PLC (it was not the case with raw distances, *e.g*. for TPM normalized with VST as revealed by their very low normalized Chi^2^ statistics; Figure 6A).

Many more co-occurring edges were generally recovered when raw CC and MI networks were compared (*e.g*. 18,334 averaged over the five pathways with MI networks vs 550 with PCC-HRR networks; Supplementary Figure 4 online). At a 1,000 vertices, all raw networks but PC contained more edges (221,297 and 85,059 in average for microarrays and TPM) than HRR-CCs networks (12,431 and 12,877). This might have resulted in more co-occurrences between MI networks. PC networks had the lowest number of co-occurring vertices (94 in average) but intersections from MI and/or raw CC had comparable vertex number (268) to intersection networks from CC-HRR (267 in average) (Supplementary Figure 4 online). These results suggest that HRR-based networks have strong overlaps. Intersections of PCC-HRR subgraphs were able to maximize the % of guide genes (mean of 75% over the 5 PLC), modularity (0.78) and normalized Chi^2^ statistic (0.70) (Figure 7). Detailed characteristics for each PLC are presented in Supplementary Figure 4 online. Modularity was generally high in the intersection between CC-HRR networks (*>*0.70) but intersections with SCC-HRR displayed lower normalized Chi^2^ values (*<*0.6). Intersection network performance in recovering GO terms was globally low (Figure 7D). The highest NV AUROCs were observed in intersections between MI networks (0.52), MI (microarrays) – raw SCC (TPM)(0.54) and raw PCC (microarrays) – raw SCC (TPM)(0.52) (Figure 7D). Intersection networks and their contents are available in Supplementary Figure 5 and Supplementary Table 4 online. Again, we found candidate genes not included in the guide gene sets that were correctly associated with other guide genes (highlighted in yellow in Supplementary Table 4 online). Taking the phenylpropanoid pathway as an example, Figure 7E shows edge and vertex co-occurrence between MI networks and Figure 7F between PCC-HRR networks. The co-occurrence network obtained from MI contained fewer guide genes (26 vs 41) and displayed lower modularity (0.49 vs 0.67) and normalized Chi^2^ statistic (0.39 vs 0.66). Although it had a higher NV AUROC (0.6 vs 0.46), its structure did not reflect that of the expected pathway (Figure 5C). For example, phenolamide related genes were not represented. Average guide gene degree (33) was below the average degree of the remaining nodes (100) indicating that guide genes were only slightly connected to other genes in this co-occurrence network from MI PLC. By contrast, guide gene degree (11.4) was very similar to the other node degree (11.1) revealing an uniform integration of guide genes with other genes in the co-occurrence network of PCC-HRR PLCs. As observed in co-occurrence in large networks (Figure 4B), RNA-seq TPM normalized with VST had slightly more edges in common with microarray networks. We therefore compared PCC-HRR PLC between microarrays and RNA-seq TPM normalized with VST. Intersection networks had very similar characteristics to that observed between microarrays and RNA-seq TPM. Although it contained slightly more co-occurring vertices and edges in average (360 and 1,252 respectively with TPM VST vs 240 and 550 with TPM), it displayed fewer guide genes (54 vs 57). TPM normalized with VST could therefore be an interesting alternative to TPM. PLC intersection networks and their description are available in Supplementary Figure 6 and Supplementary Table 5 online.

**Figure 7.**
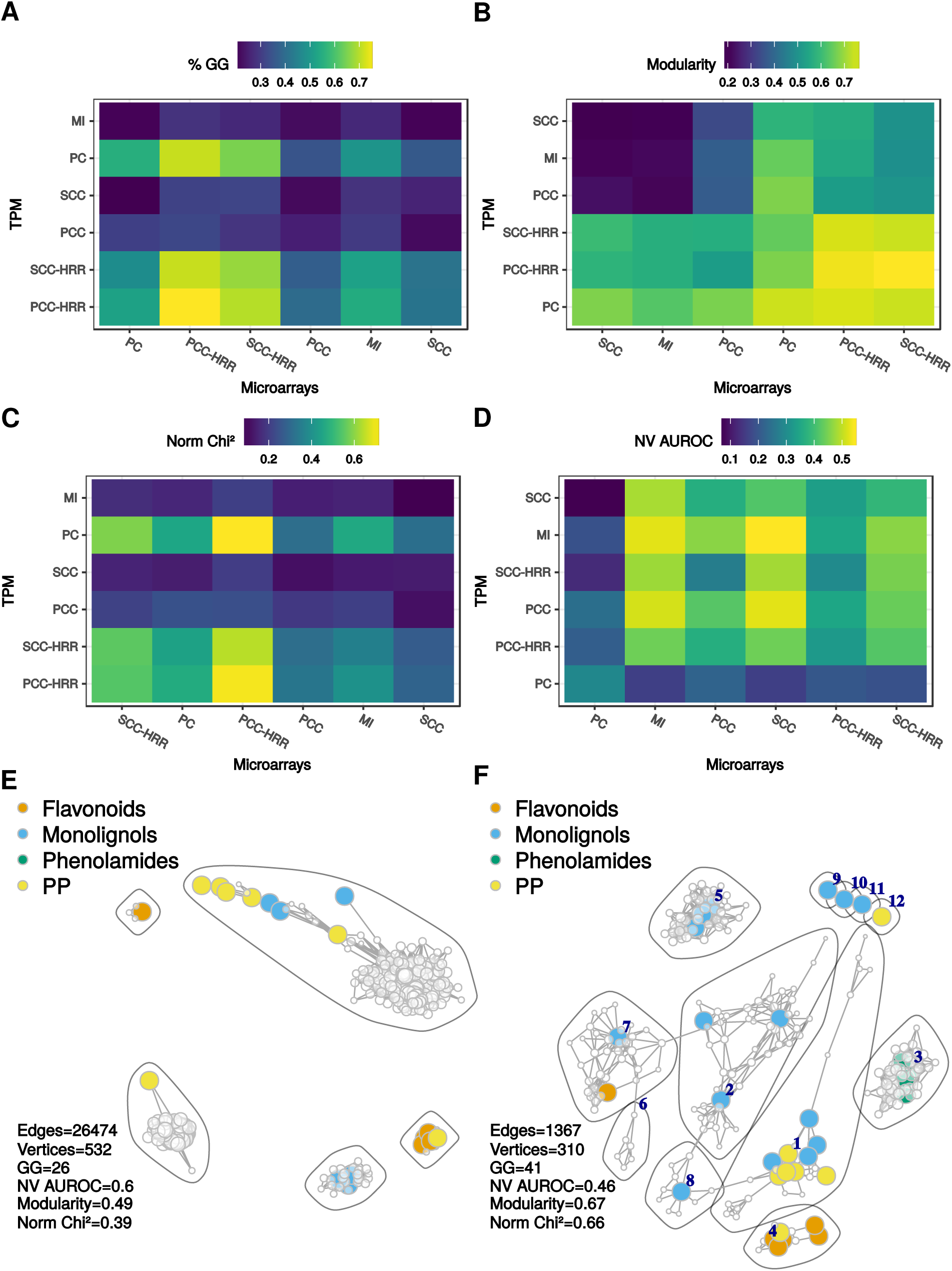
Characteristics of co-occurrence networks between microarrays and RNA-seq TPM. Percentage of guide genes (GG; A), modularity (B), normalized Chi^2^ statistic (agreement with guide gene partitioning, C) and NV AUROC (GO term performance, D) were averaged over the 5 PLCs. Labels are ordered according to a hierarchical clustering. Co-occurrence networks obtained from phenylpropanoid PLC obtained with MI (E) or PCC-HRR (F). GG corresponds to guide gene number in the networks. Community numbers in F are indicated in deep blue and can be used to access Supplementary Table 4 online.

## Discussion

Pathway Level-Correlation (PLC) is an interesting approach to capture biologically relevant transcriptional relationships using guide genes (e.g. genes involved in a same metabolic pathway) from transcriptome-wide co-expression networks. Our present work highlights that distances between genes calculated with highest reciprocally ranked PCC (PCC-HRR) improve PLC. The main improvement was guide gene representation. PCC-HRR based PLCs contained more guide genes than observed with other distances and they were generally more correctly partitioned into expected sub-pathways in the co-expression network. This was associated with a lower mean node degree and a higher modularity but also with a slightly weaker performance in GO term recovery. Our results propose that modularity and normalized Chi^2^ values could be used as reliable indicators of PLC quality. We also observed that edge and vertex co-occurrences in PLCs obtained with PCC-HRR and microarray and RNA-seq TPM data can be used to construct relevant networks. A surprising observation was that in our conditions, for most combinations tested, true positive rates remained higher than false positive rates in spite of increasing network sizes. A similar trend using small *E. coli* and *S. cerevisiae* networks (*<*110 nodes) has been previously observed with CCs^7^. This suggests that co-expression studies should test different confidence thresholds to efficiently capture gene associations. Evaluating network quality was done in respect of the Arabidopsis reference GO annotation set. We found that the NV AUROC^14^ evaluates networks efficiently and was generally in accordance with significantly enriched GO term counts and TPR vs FPR curves. NV AUROC has the advantage of being a more global measure of predictability (values above 0.6 can be considered as moderate). Different distance measurements displayed different efficiencies according to the dataset but as a general trend, performance of the different combinations were similar (e.g. between microarrays and RNA-seq TPM in Figure 3B). The same performance was obtained for different topologies: high node degree (more edges and fewer vertices) for MI and raw CC networks vs lower node degree (fewer edges and more vertices) for CC-HRR networks. PC networks displayed a high performance with microarray data only, suggesting that PCs calculated with ‘corpcor’ R package may not be recommended for RNA-seq data. A recent study has focused on metabolic pathways in plants using mutual ranks, another CC ranking method^12^. Complementary to this previous work, we found that ranking CCs increases vertex number without penalizing absolute network performance. Contrastingly, an opposite trend was observed in another study^24^, where larger networks displayed a lower Matthew Coefficient when compared to protein-protein interactions or regulatory networks. This indicates that different absolute performance measurements lead to different results and interpretations but this might also be due to our datasets which were larger than theirs. Another advantage of CC-HRR was that it clearly homogenized network characteristics from differently normalized RNA-seq datasets in addition to increase the number of co-occurring edges between microarrays and RNA-seq.

As revealed recently^25^, highlighting correlations between genes may require specific data processing or distance algorithms best suited to their query pathway. We also found that each of the five PLCs performed best with specific RNA-seq normalizations (Figure 6C & D) but RNA-seq TPM processed with PCC-HRR always provided informative networks which can be used as reliable starting point because they matched well expected pathway structure. In our case, the different data normalizations had a relatively weak effect on PLC characteristics especially when CCs were used with HRR. In a comparative analysis^24^, the authors have shown that PCC networks from VST normalized counts were more comparable to those from microarrays. In our case, VST normalization slightly improved the overlap between RNA-seq TPM and microarrays both at the global and targeted levels. This normalization can thus be further considered for co-expression studies. A fast greedy approach maximizing modularity was used to detect communities within PLC subgraphs. Guide gene partitioning in these communities was compared to expected partitions in subpathways with a normalized Chi^2^ test (Figure 5A). We found that correct guide gene partitioning was negatively correlated with NV AUROCs but positively with modularity. Subnetworks with highest NV AUROCs but lower modularity such as those obtained with MI represented fewer guide genes and displayed large edge numbers. In these networks, guide genes formed inappropriate structures (Supplementary Figure 5 online). We applied PLC to five pathways varying in size and nature. For the four metabolic pathways, PLC extracted from PCC-HRR based networks were able to cluster guide genes in the proper subpathways (Supplementary Figure 5 online). Guide genes were associated in communities resembling subpathways and containing genes not included in the query gene set but known to be involved with the given pathway or being good candidates to be functionally validated (Supplementary Table 4 online). A similar PLC approach has been recently performed^12^ using the Arabidopsis aliphatic glucosinolate pathway. The authors have successfully reconstructed this pathway and identified a new candidate glucosyltransferase that could be part of it. This demonstrated again that PLC is a powerful approach to complete biological pathways. When tested with a signaling pathway, we found that PLCs also displayed meaningful communities. For example, the CK signaling pathway is physiologically well known but its organization at the molecular level is far from being understood^21^. In particular, it is unclear how multi-family members of each signaling level (receptor, transducer and response regulator) interact with each other to drive a specific physiological response. In the PLC dedicated to the CK signaling pathway, PCC-HRR with microarrays suggested preferential transcriptional associations that have been described in the literature [36]. For example AHP2, AHP3 and AHP5 were grouped in the same module (module 7 Supplementary Figure 5 and Supplementary Table 4 online). These three AHPs have been reported to negatively regulate tolerance to abiotic stress [40]. The same community also contained AHK3, ARR1 and ARR2. Those three members are known to regulate primary root meristem activity and senescence^26^. AHK4, AHK2 and ARR14 which have been shown to regulate shoot apical meristem activity were grouped in the same community^27^. In addition, we saw clear associations between ETR1 and AHK3 in individual PLC subgraphs. Such association highlights crosstalk already known between CK and ethylene signaling pathways^22^. The co-occurrence pathway was relatively sparse in contrast to the metabolic pathways (Supplementary Figure 5 online). It is possible that vertex number for this analysis (1,000) might have been too small to capture complex associations within this signaling pathway. Using VST normalized TPM increased edge and vertex number in the co-occurrence network (Supplementary Figure 6 and Supplementary Table 5 online). The above-described associations were also found in this co-occurrence network. While effective in revealing strong gene associations, merging PLC from microarray and RNA-seq data could miss other relevant associations. First, experimental conditions represented by each starting dataset are not completely overlapping. Together with inherent differences due to dynamic range, this leads to networks with very different edge compositions and node degrees^14^, explaining the relative weak overlap between networks. Second, RNA-seq expression data include genes that are not included in the GPL198 microarray. As an example, some important genes in aliphatic glucosinolate biosynthesis were not represented in a previous Arabidopsis microarray dataset but found in RNA-seq expression matrices from other related species^12^.

To capture transcriptional environment of a query gene list, distance calculations have to be performed on the whole transcriptome. Calculating partial correlations was particularly challenging but using a covariance shrinkage estimator worked well in terms of computing performance. It took less than 2h for RNA-seq expression matrices but more than 12h for the microarray dataset. By contrast, our program which is freely available at (https://github.com/EA2106-Universite-Francois-Rabelai-Expression-network-analysis) was able to calculate PCC-HRR in less than 3h for both datasets. As PCC-HRR resulted in relevant networks, this tool can be useful for further studies requiring many computations such as analyzing sample size impact on PLC or testing other normalization methods.

The present work demonstrates that Pearson’s Correlation Coefficients (PCC) on which highest reciprocal ranking (HRR) was applied can be used to construct reliable global and targeted networks. When considering Pathway Level Correlation (PLC) with a set of guide genes, three reliable measures can be used for evaluation: NV AUROC as a global indicator of GO recovery (expecting values*>*0.5), modularity (between 0 and 1, 1 being the best network partition) and normalized Chi statistic (between 0 and 1, 1 indicating a perfect match with an expected partition). Clustering guide genes correctly was at the expense of capturing GO terms and dataset x distance measurement combination should be carefully selected to construct reliable PLC. Although specific RNA-seq data normalizations may be adapted to each pathway of interest, using TPM with PCC-HRR generated accurate and safe PLC. Using PCC with HRR also increased the quality of co-ocurrence networks between RNA-seq and microarrays.

## Methods

### Microarray data preparation

Experiment accessions (GSE) for GPL198 (Arabidopsis ATH1, 22,746 genes) were retrieved from ArrayExpress (Supplementary Table 6 online). Signal intensities per probe were generated with R [16]using the ‘arrayexpress’ package^28^. The function ‘getAE’ was used to convert the raw signal CEL files. Array normalization was performed per GSE using the ‘justRMA’ function of the ‘affy’ package. This procedure applies a background correction together with a quantile normalization to correct for biases within arrays and finally returns log2-transformed corrected signal intensities. All 10,095 arrays were combined into a single file and subjected to a quality control based on upper quartile dispersion (75%) and Kolmogorov-Smirnov statistical testing for outliers using an empirical cumulative distribution function as described previously^29^. A total of 142 arrays were considered outliers in the two tests and discarded from the final matrix. Each array was finally centered and scaled individually.

### RNA-seq data preparation

2,549 RNA-seq accessions obtained for *A. thaliana* were retrieved from ArrayExpress. Fastq files were obtained from the SRA after converting .sra files with the SRA ToolKit function ‘fastq-dump’ with the –split-files option for paired-end sequencing runs. Reads were systematically trimmed with Trimmomatic using adapter files according to the Illumina platform used for the runs (ref). Trimmed reads were pseudo-aligned to predicted transcripts from the representative gene models of Arabidopsis TAIR genome v10 (33,604 transcripts) with Salmon v0.7.2 using the variational Bayesian EM algorithm mode to improve abundance estimation^30^. Only samples displaying a mapping rate of reads *>*30% were kept, resulting in a final matrix containing 1,676 samples (Supplementary Table 6 online). RNA-seq counts were used as non-normalized raw counts or expressed as Transcript per Million to correct for sequencing depth. Normalization by Variance Stabilizing Transformation (VST) was performed with the DESeq2 R package. This normalization method aims at limiting the variance dependence to the mean^31^.

### Distance calculations

Before calculations, zero-variance genes were discarded. CCs (Pearson or Spearman) are computationally intensive particularly in the case of large matrices. Highest Reciprocal Ranking (HRR) of CCs for genes A and B is calculated as max(rank(CC(A,B)), rank(CC(B,A)). For each gene, all CC values are first transformed as ranks, with 0 corresponding to the gene rank against itself. Ranks are subsequently compared and the highest value is retained for each gene pair. We developed a tool written in C allowing the easy parallelization of these computations. Briefly, for a given initial matrix containing *n* genes and *p* samples, the number of cores *c* allocated is used to split the dataset into *n*/*c* submatrices. In case of non-integer value, the last line of the matrix is replicated (without incidence on PCC or rank values) so that *n*/*c* is an integer. PCC or HRR are then calculated for each gene pair using communication between CPUs with Message Passing Interface. The program delivers c files containing *n*/*c* x *n* values corresponding to PCC or HRR. This program is freely available on Github (https://github.com/EA2106-Universite-Francois-Rabelais/Expression-network-analysis). To calculate SCCs, expression values were first ranked in R. Mutual information (MI) which is reported to better capture non linear relationships^8^ were calculated with the ‘knni.all’ function of the Parmigene R package^32^. This function estimates MI using a *k*-nearest neighbor. Partial correlations were challenging to compute on genome scale expression matrices. Partial correlations are usually calculated from multiple linear regressions or by inverting the correlation matrix and used in Graphical Gaussian Models^33^. Our expression matrices had many more variables (genes) than samples therefore regression methods would have required a Lasso or Ridge penalization to estimate coefficients. However, this procedure generally leads to memory errors when considering more than 30,000 variables. We found that the most computationally appropriate method in our case was to estimate shrinkages of partial correlations with the R package ‘corpcor’ (http://strimmerlab.org/software/corpcor/). This package is maintained by Korbinian Strimmer’s team^34, 35^. We used ‘pcor.shrink’ function which relies on the inversion of the shrunken estimated covariance matrix to estimate partial correlations and which is suited for matrices with more genes than samples.

### Reference dataset

We used the Arabidopsis Gene Ontology (GO) standard dataset to assess network quality. The annotation file provided by the AGRIGO database^36^ and was filtered out to remove all terms with a IEA evidence code and keep only functionally attributed terms. We also removed GO terms represented by less than 5 genes or more than 100 to remove non-specific terms.

### Global Network analysis

Construction: For each dataset x distance combination, we dynamically set a threshold to obtain arbitrary lists of 10 million best gene pairs (with CC above or HRR below that threshold), *i.e*. less than 2% of the total possible edges. Networks were then constructed with the 1, 5, 10, 20, 40, 60 or 80% best pairs from these lists. Thresholds used to get the 10 million gene pairs are reported in S2 Table. Global networks were analyzed as adjacency matrices in R. Network characteristics: besides classical topological characteristics such as vertex and edge numbers and mean node degree (the average number of connections for each vertex), we evaluated network quality by comparison with the reference dataset (Figure 2). In a first approach, we built a confusion matrix by classifying edges as false or true positives, considering edges as valid if both genes were annotated with at least one same GO term. In this confusion matrix, true positives (TP) corresponded to gene pairs also found in the GO annotation, false positive (FP) to genes associated in the network but not in the GO annotation, false negatives (FN) to pairs in the GO annotation not predicted in the network and finally true negatives (TN) genes pairs not predicted in the network and the annotation table. This confusion matrix was used to calculate True Positive Rates (TPR) and False Positive Rates (FPR). TPR and FPR were obtained at various confidence thresholds (i.e. for networks differing in sizes) and used to draw a TPR vs FPR curve as described elsewhere^11^. These curves were only partial because we included only the first 10 million best pairs. This was useful to pinpoint the importance of low FPR^37^. In the second and third approaches, we relied on the guilt-by-association principle to estimate network predictability. In the second method, we used the ‘predictions’ function of EGAD R package^38^. For each gene, this function counts the number of connected genes annotated with an identical GO term and divides this count by the gene’s degree. These scores are next ordered decreasingly to construct a TPR vs FPR curve for each network. It differs from the first approach described above because here TPR and FPR are not obtained from different confidence thresholds (and from different networks) but from all possible true positive and false positive edges in the current network. A global Area Under Receiver Operating Characteristic (global AUROC) was calculated from each of these TPR/FPR curves. In the third method, predictability was evaluated using a neighbor voting (NV) algorithm. In this case, an AUROC is calculated for each GO term from the ability of genes to predict the GO annotation of their direct neighbors in a 3-fold cross-validation^14, 39^. A mean NV AUROC was calculated for each network. In addition to ROC analysis, we counted GO terms that were significantly enriched with gene pairs using a hypergeometric test with R.

### Pathway Level Correlation

Construction: In PLC, subnetworks were constructed from global networks (see above) by keeping edges connecting at least one guide gene. Guide gene lists are indicated in Supplementary Table 2 online. The R package ‘igraph’^40^ v1.0.1 was used to construct and visualize these targeted networks with a force-directed layout (Fruchterman-Reingold). Community Detection: Modules containing densely connected vertices were estimated within each network by using a fast greedy approach which aims at maximizing modularity of the detected communities^23^. Modularity measures how good a network partition is by calculating for each gene the number of edges within its community against its total node degree. The fast greedy approach optimizes modularity over all possible divisions of the network and has been shown to perform well on large networks. Guide genes clustering within the communities was compared to expected partitions in sub-pathway with a Pearson’s Chi^2^ test and Monte-Carlo simulated *p*-values with 2,000 replicates. This test was based on a contingency table with dimensions *n* x *m* (*n*, sub-pathway number, *m* community number in the co-expression network) and each entry corresponding to the number of genes being in communities n_i_ and m_j_, with i=1 to *n* and j=1 to *m*. Because Chi^2^ statistic depends on sample number, values were normalized by dividing them to the maximal expected value (the ideal partition) of each pathway. This resulted in a score ranging from 0 to 1, 0 being a random distribution of guide genes in the network and 1 to the exact partitioning.

## Acknowledgements

We deeply acknowledge the Fédération CaSciModOT (CCSC Orléans-Tours, France), Jean-Louis Rouet and Laurent Catherine for help and access to the Région Centre computing grid. We also thanks Yann Jullian for access and help on University computer resources. This study was supported by the Région Centre-Val de Loire, France (SiSCyLi grant). Doctoral Fellow attributed to F.L. and D.D. was jointly funded by the Région Centre-Val de Loire, France and the Ministère de l’Enseignement Supérieur et de la Recherche, France.

## Author contributions statement

F.L., O.P., J.C., N.G. and T.D.D.B. conceived the experiment(s), F.L., D.D., O.P., M.C., S.B., V.C. and R.D.D.B conducted the experiment(s), F.L., S.B., G.G., J.C., J.O.C. and T.D.D.B. analyzed the results. All authors reviewed the manuscript.

## Data availability

All datasets generated during and/or analyzed during the current study are available from the corresponding author on reasonable request.

## Supplementary Information

**Supplementary Figure 1: Network properties in dataset-distance measurement combinations.** Global network characteristics (Number of significantly enriched GO terms, global and NV AUROCs) were expressed in function of vertex or edge number. The horizontal dashed line indicates a 0.6 AUROC value taken as an arbitrary threshold separating good and poor network predictability. For each dataset, TPR=f(FPR) curves are also presented with dashed line corresponding to a random selection (with AUROC *<*0.5). These curves are partial and the max FPR values were obtained for 10 million gene pairs.

**Supplementary Table 1: Threshold values to get 10 million best gene pairs**.

**Supplementary Figure 2: Workflow for Pathway-Level Correlation. Lists of best co-expressed genes are established for each guide (or bait) gene.** Redundancies among these lists (associated genes) connect guide genes to construct the PLC network. Terms ‘guide gene’ and ‘associated genes’ have been introduced by Lisso et al^2^.

**Supplementary Table 2: Guide gene accessions**.

**Supplementary Figure 3. PLC subgraphs for the carbohydrate (A), fatty acid (B), terpene (C) and cytokinin (D) pathways.** For each PLC, the expected partitioning of guide genes is indicated in the left panel and is compared to PLC subgraphs with higher predictability and lower modularity (center; calculated with MI) or PLC subgraphs with lower predictability and higher modularity (right; calculated with PCC-HRR). Colored vertices correspond to genes encoding enzymes catalyzing steps of similar color in the expected pathway. A and B were drawn from RNA-seq TPM networks while C and D from microarray networks. Community numbers in PCC-HRR networks are indicated in deep blue and can be used to access Supplementary Table 3 online. Polygons surrounding vertices delimit communities.

**Supplementary Table 3: Gene lists from PLC obtained with PCC-HRR Genes highlighted in yellow correspond to non-guide genes but known to be involved in the pathway**.

**Supplementary Figure 4: PLC based on microarray and TPM data.** Subgraphs were constructed with the 6 distance measurements (MI, PC, raw PCC, raw SCC, PCC-HRR and SCC-HRR) and aligned to find co-occurring edges and vertices. (A) Number of co-occurring vertices and edges. The first distance in each label correspond to microarrays and the second to TPM. Points are half-colored according to the ranking applied to the initial distance. For each intersection graph, % of guide genes (B), normalized Chi^2^ statistic (agreement with expected guide gene partitioning, C), modularity (D) and NV AUROC (GO recovery performance, E) were calculated.

**Supplementary Figure 5: Co-occurrence networks from PCC-HRR PLC constructed with microarrays and RNA-seq TPM**. Community numbers are indicated in deep blue and can be used to access Supplementary Table 4 online.

**Supplementary Table 4: Gene lists from co-occurrence networks between PCC-HRR PLC obtained with microar-rays and RNA-seq TPM.** Genes highlighted in yellow correspond to non-guide genes but known to be involved in the pathway.

**Supplementary Figure 6: Co-occurrence networks from PCC-HRR PLC constructed with microarrays and RNA-seq TPM normalized with VST.** Community numbers are indicated in deep blue and can be used to access Supplementary Table 5 online.

**Supplementary Table 5: Gene lists from co-occurrence networks between PCC-HRR PLC obtained with microar-rays and RNA-seq TPM normalized with VST**.

**Supplementary Table 6: Microarray and RNA-seq accessions used in this study**.

## References

1. Oliver, S. Proteomics: guilt-by-association goes global. Nat. 403, 601–603 (2000).

2. Lisso, J., Steinhauser, D., Altmann, T., Kopka, J. & Müssig, C. Identification of brassinosteroid-related genes by means of transcript co-response analyses. Nucleic Acids Res. 33, 2685–2696 (2005).

3. Wei, H. et al. Transcriptional coordination of the metabolic network in arabidopsis. Plant physiology 142, 762–774 (2006).

4. Ruiz-Sola, M. Á. et al. Arabidopsis geranylgeranyl diphosphate synthase 11 is a hub isozyme required for the production of most photosynthesis-related isoprenoids. New Phytol. 209, 252–264 (2016).

5. Guerin, C. et al. Gene coexpression network analysis of oil biosynthesis in an interspecific backcross of oil palm. The Plant J. 87, 423–441 (2016).

6. Coman, D., Rütimann, P. & Gruissem, W. A flexible protocol for targeted gene co-expression network analysis. Plant Isoprenoids: Methods Protoc. 285–299 (2014).

7. Maetschke, S. R., Madhamshettiwar, P. B., Davis, M. J. & Ragan, M. A. Supervised, semi-supervised and unsupervised inference of gene regulatory networks. Briefings bioinformatics 15, 195–211 (2013).

8. de Siqueira Santos, S., Takahashi, D. Y., Nakata, A. & Fujita, A. A comparative study of statistical methods used to identify dependencies between gene expression signals. Briefings bioinformatics 15, 906–918 (2013).

9. Li, Y., Pearl, S. A. & Jackson, S. A. Gene networks in plant biology: approaches in reconstruction and analysis. Trends plant science 20, 664–675 (2015).

10. Serin, E. A., Nijveen, H., Hilhorst, H. W. & Ligterink, W. Learning from co-expression networks: possibilities and challenges. Front. plant science 7 (2016).

11. Obayashi, T. & Kinoshita, K. Rank of correlation coefficient as a comparable measure for biological significance of gene coexpression. DNA research 16, 249–260 (2009).

12. Wisecaver, J. H. et al. A global co-expression network approach for connecting genes to specialized metabolic pathways in plants. The Plant Cell Online tpc–00009 (2017).

13. Mutwil, M. et al. Assembly of an interactive correlation network for the arabidopsis genome using a novel heuristic clustering algorithm. Plant Physiol. 152, 29–43 (2010).

14. Ballouz, S., Verleyen, W. & Gillis, J. Guidance for rna-seq co-expression network construction and analysis: safety in numbers. Bioinforma. 31, 2123–2130 (2015).

15. Song, L., Langfelder, P. & Horvath, S. Comparison of co-expression measures: mutual information, correlation, and model based indices. BMC bioinformatics 13, 328 (2012).

16. Besseau, S. et al. Flavonoid accumulation in arabidopsis repressed in lignin synthesis affects auxin transport and plant growth. The Plant Cell 19, 148–162 (2007).

17. Zhang, Y. et al. Phenolic compositions and antioxidant capacities of chinese wild mandarin (citrus reticulata blanco) fruits. Food chemistry 145, 674–680 (2014).

18. Winkel-Shirley, B. Flavonoid biosynthesis. a colorful model for genetics, biochemistry, cell biology, and biotechnology. Plant physiology 126, 485–493 (2001).

19. Elejalde-Palmett, C. et al. Characterization of a spermidine hydroxycinnamoyltransferase in malus domestica highlights the evolutionary conservation of trihydroxycinnamoyl spermidines in pollen coat of core eudicotyledons. J. experimental botany 66, 7271–7285 (2015).

20. Kanehisa, M., Sato, Y., Kawashima, M., Furumichi, M. & Tanabe, M. Kegg as a reference resource for gene and protein annotation. Nucleic acids research 44, D457–D462 (2016).

21. Hwang, I., Sheen, J. & Müller, B. Cytokinin signaling networks. Annu. review plant biology 63, 353–380 (2012).

22. Zdarska, M. et al. Illuminating light, cytokinin, and ethylene signalling crosstalk in plant development. J. experimental botany 66, 4913–4931 (2015).

23. Clauset, A., Newman, M. E. & Moore, C.. Finding community structure in very large networks. Phys. review E 70, 066111 (2004).

24. Giorgi, F. M., Del Fabbro, C. & Licausi, F. Comparative study of rna-seq-and microarray-derived coexpression networks in arabidopsis thaliana. Bioinforma. 29, 717–724 (2013).

25. Uygun, S., Peng, C., Lehti-Shiu, M. D., Last, R. L. & Shiu, S.-H. Utility and limitations of using gene expression data to identify functional associations. PLoS computational biology 12, e1005244 (2016).

26. Jiang, L. et al. Strigolactones spatially influence lateral root development through the cytokinin signaling network. J. experimental botany 67, 379–389 (2015).

27. Wang, L. & Chong, K. The essential role of cytokinin signaling in root apical meristem formation during somatic embryogenesis. Front. plant science 6 (2015).

28. Kauffmann, A. et al. Importing arrayexpress datasets into r/bioconductor. Bioinforma. 25, 2092–2094 (2009).

29. Feltus, F. A., Ficklin, S. P., Gibson, S. M. & Smith, M. C. Maximizing capture of gene co-expression relationships through pre-clustering of input expression samples: an arabidopsis case study. BMC systems biology 7, 44 (2013).

30. Patro, R., Duggal, G., Love, M. I., Irizarry, R. A. & Kingsford, C. Salmon provides fast and bias-aware quantification of transcript expression. Nat. Methods 14, 417–419 (2017).

31. Love, M. I., Huber, W. & Anders, S. Moderated estimation of fold change and dispersion for rna-seq data with deseq2. Genome biology 15, 550 (2014).

32. Sales, G. & Romualdi, C. parmigene—a parallel r package for mutual information estimation and gene network reconstruction. Bioinforma. 27, 1876–1877 (2011).

33. López-Kleine, L., Leal, L. & López, C. Biostatistical approaches for the reconstruction of gene co-expression networks based on transcriptomic data. Briefings functional genomics 12, 457–467 (2013).

34. Schäfer, J. & Strimmer, K. Learning large-scale graphical gaussian models from genomic data. In AIP Conference Proceedings, vol. 776, 263–276 (AIP, 2005).

35. Schaefer, J., Opgen-Rhein, R. & Strimmer, K. corpcor: efficient estimation of covariance and (partial) correlation. r package version 1.4. 7 (2007).

36. Du, Z., Zhou, X., Ling, Y., Zhang, Z. & Su, Z. agrigo: a go analysis toolkit for the agricultural community. Nucleic acids research 38, W64–W70 (2010).

37. Schrynemackers, M., Küffner, R. & Geurts, P. On protocols and measures for the validation of supervised methods for the inference of biological networks. Front. genetics 4 (2013).

38. Ballouz, S., Weber, M., Pavlidis, P. & Gillis, J. Egad: ultra-fast functional analysis of gene networks. Bioinforma. 33, 612–614 (2016).

39. Gillis, J. & Pavlidis, P. The impact of multifunctional genes on” guilt by association” analysis. PloS one 6, e17258 (2011).

40. Csardi, G. & Nepusz, T. The igraph software package for complex network research. InterJournal, Complex Syst. 1695, 1–9 (2006).

